# Magic pools: parallel assessment of transposon delivery vectors in bacteria

**DOI:** 10.1101/158840

**Authors:** Hualan Liu, Morgan N. Price, Robert Jordan Waters, Jayashree Ray, Hans K. Carlson, Jacob S. Lamson, Romy Chakraborty, Adam P. Arkin, Adam M. Deutschbauer

**Author notes:** Supplementary website for paper: http://genomics.lbl.gov/supplemental/magicpools/.

## Abstract

Transposon mutagenesis coupled to next-generation sequencing (TnSeq) is a powerful approach for discovering the functions of bacterial genes. However, the development of a suitable TnSeq strategy for a given bacterium can be costly and time-consuming. To meet this challenge, we describe a parts-based strategy for constructing libraries of hundreds of transposon delivery vectors, which we term “magic pools”. Within a magic pool, each transposon vector has a different combination of promoters and antibiotic resistance markers as well as a random DNA barcode sequence, which allows the tracking of each vector during mutagenesis experiments. To identify an efficient vector for a given bacterium, we mutagenize it with a magic pool and sequence the resulting insertions; we then use the best vector to generate a large mutant library. We used the magic pool strategy to construct transposon mutant libraries in five genera of bacteria, including three genera of the phylum *Bacteroidetes*.

## Introduction

Transposons are mobile genetic elements that can translocate from a donor site to one of many target sites, without any homology requirement (1). Since transposons were first identified in maize over sixty years ago, they have been widely used to introduce insertional mutations for gene function identification, both in prokaryotes and eukaryotes. Among various transposons, Tn5 and *mariner* have been commonly used in mutagenesis of bacterial genomes (2) due to their low target site bias and high degree of randomness (3, 4).

During the last decade, transposon mutagenesis has been coupled to next-generation sequencing to dramatically advance gene function discovery in bacteria (5-7). In these strategies, a large population of transposon insertion mutants can be pooled together and the relative abundance of all of the mutant strains can be monitored by next-generation sequencing of the genomic DNA flanking the transposon insertion. By constructing very large libraries of mutants containing thousands to millions of unique insertion strains, the genome-wide interrogation of mutant fitness and hence gene fitness can be conducted in parallel in a single tube. The first such strategy, termed TnSeq, was used in *Streptococcus pneumoniae* to globally measure single gene fitness, and also to screen genetic interactions (8). After this initial report, a number of TnSeq-like variants have been described including TraDIS (9), HITS (10), IN-Seq (11), or TnLE-seq (12).

Our group has recently developed a variant of TnSeq termed random barcode transposon site sequencing (RB-TnSeq) that simplifies genome-wide transposon fitness assays in bacteria (13). In RB-TnSeq, random DNA barcodes are incorporated into a transposon delivery vector, and only one initial round of TnSeq is needed to map the insertion site of the transposon mutant to the unique barcode that identifies that specific mutant. To quantify strain abundance and calculate gene fitness scores, one only needs to PCR amplify and sequence the DNA barcodes (BarSeq (14)). BarSeq is a simple assay and 96 samples can be multiplexed on a single lane of Illumina HiSeq (13). Therefore, RB-TnSeq greatly accelerates functional genomics in bacteria with established genetic systems. However, for any given bacterium, it is likely that neither of the two barcoded transposon delivery vectors described in the original work will function. For example, both vectors used a kanamycin resistance gene as a selectable antibiotic marker, which precludes the mutagenesis of naturally kanamycin-resistant bacteria. More broadly, the development of a functional genetic system in any new bacterium can be time-consuming, with multiple cycles of trial and error before a functioning system is found.

Here we describe an approach for testing hundreds of transposon delivery vectors in parallel. We constructed magic pools of many different transposon delivery vectors, each with a unique DNA barcode inside the transposon. These magic pools can be constructed efficiently using a parts-based strategy and Golden Gate assembly. The sequence of each vector, including its associated DNA barcode, can be determined by long-read DNA sequencing. Given a magic pool and a target bacterium, we can assess the efficiency of hundreds of different transposon vectors in parallel by performing a single mutagenesis experiment followed by TnSeq. Once the best vector has been identified, it can be rapidly reconstructed, barcoded, and used to build a high-coverage mutant library for RB-TnSeq. To demonstrate this approach, we built four magic pools with Tn5 or *mariner* transposase and with kanamycin or erythromycin as the selectable antibiotic marker. Using these magic pools, we were able to rapidly generate high-coverage transposon mutant libraries for five different genera of bacteria, including three genera of Bacteroidetes.

## Results

### Overview of the magic pool approach

Our approach is illustrated in Figure 1. First, we split a traditional transposon delivery vector into five separate parts that can be readily reassembled using Golden Gate assembly (15, 16). Part1 contains the majority of the open reading frame (ORF) for either the Tn5 or the *mariner* transposase and one copy of its inverted repeat (13); part2 is the promoter for driving the expression of the antibiotic selection marker; part3 is the ORF of the antibiotic selection marker; part4 contains an ampicillin drug cassette for selection during cloning, *oriT* for the initiation of conjugation, a conditional R6K origin of replication, the second copy of the inverted repeat, and optionally a random 20-nucleotide DNA barcode; and part5 is the promoter for and a short region of the 5’ end of the transposase. In this work, we refer to a “promoter” as the entire upstream region of the ORF, which therefore includes both the promoter and the ribosome binding site. The promoter parts are either from known antibiotic resistance cassettes, or based on the sequences upstream of predicted essential genes from diverse bacteria (Materials and Methods). Parts 1, 4, and 5 contain sequences that are unique to either Tn5 or *mariner* transposon systems, either the transposon inverted repeats or the transposase. Therefore, these parts are sequence specific for their transposon type. The same parts 2 and 3 are used by both Tn5 and *mariner* magic pools.

**Figure 1.**
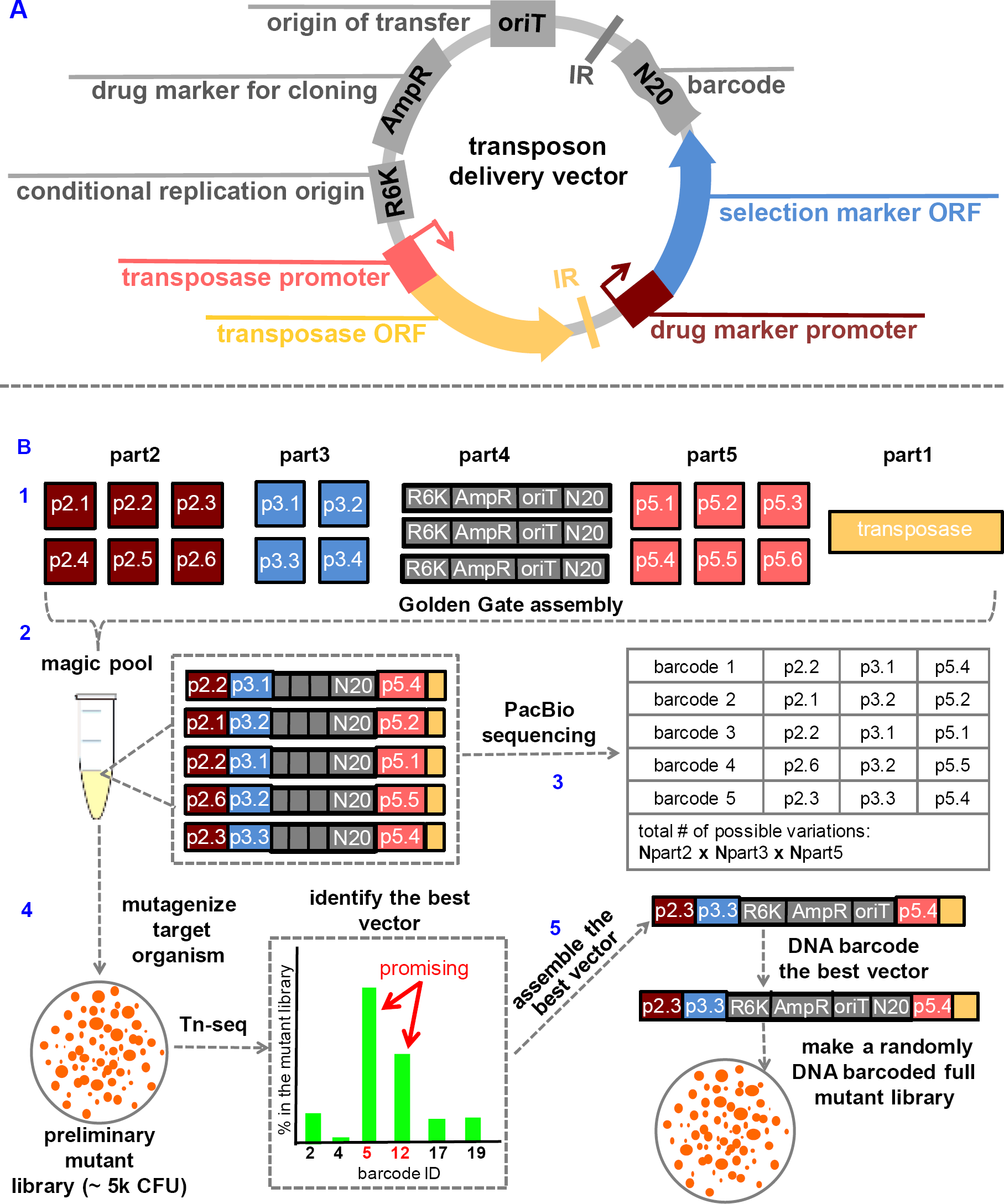
Overview of the magic pool strategy. (A) Basic structure of a typical transposon delivery vector (not drawn to scale). IR is an inverted repeat for the specific transposase. We dissected the transposon delivery vector into 5 different parts compatible with Golden Gate assembly, and each color represent a part. (B) General workflow of construction and application of magic pools. (1) Variants of the five different parts are designed, cloned into a part-holding vector, confirmed by sequencing, and archived. (2) The parts vectors are mixed and assembled using Golden Gate assembly to produce the magic pools of transposon delivery vectors. (3) The magic pool vectors are characterized by DNA sequencing whereby each unique DNA barcode (N20) is linked to a specific combination of parts. (4) Preliminary mutant libraries are made using the magic pool and TnSeq is performed to link the DNA barcode to insertion site thereby simultaneously assessing the efficacy of the vectors in the magic pool. (5) The best vector is reassembled using the archived parts, fully barcoded with millions of random DNA barcodes, and a full RB-TnSeq transposon mutant library is constructed. ColE1: replication origin colE1; oriT: origin of transfer; ampR: beta-lactam resistance cassette; R6K: conditional replication origin; N20: random 20 nucleotide DNA barcode.

A combinatorial pool of DNA-barcoded transposon vectors can be constructed by Golden Gate assembly with part1, a library of promoters (part2 and part5) and antibiotic selection markers (part3), and a randomly barcoded part4. This magic pool of transposon delivery vectors is characterized by PacBio sequencing, which provides sequencing reads that are long enough to cover all of the parts and the random DNA barcode. We also used Illumina sequencing of the barcodes (BarSeq (14)) to ensure that they are accurate.

Given a bacterium of interest, we make small, preliminary mutant libraries with 1,000-5,000 mutants using the magic pools. In practice, we usually try both a *mariner* magic pool and a Tn5 magic pool. We use DNA sequencing to identify the most effective transposon delivery vector for that particular bacterium. To determine which parts give the largest number of mutants, it is sufficient to just amplify and sequence the barcodes with BarSeq. However, transposons can have uneven insertion distribution across the chromosome or exhibit insertion strand bias, so the parts that result in the most insertions may not actually be the best for constructing the full mutant library. Therefore, we also perform TnSeq to sequence the transposon insertion junction and link the location of each insertion in the genome to the barcode, and hence to the parts.

Once an optimal transposon delivery vector is identified from the magic pool, we assemble that specific vector (with non-barcoded part4) using the archived parts collection, incorporate millions of random DNA barcodes using another round of Golden Gate cloning, and construct the full barcoded transposon mutant library. In the magic pool, the purpose of the barcode is to identify the parts of the transposon delivery vector, but in the final library, the purpose of the barcode is to simplify the quantification of strain abundance in a genome-wide fitness assay with BarSeq (13).

### Proof of concept with kanamycin resistance marker

As a proof of principle, we constructed two small magic pools using kanamycin as the antibiotic selection marker and either Tn5 or *mariner* transposase. The *mariner*-Kan magic pool is based on the original RB-TnSeq *mariner* vector pKMW3 (13). We constructed a pool of 10 defined *mariner* transposon delivery vector designs (pTGG31-pTGG40; **Table S4**) by including 2 different promoters for the kanamycin selection marker (part2), 5 different promoters for the *mariner* transposase (part5), and a randomly DNA barcoded part4. We kept the original kanamycin selection marker ORF (part3) and the *mariner* transposase ORF (part1) from pKMW3 constant in all 10 vector designs. The promoters for part5 are from predicted essential genes in Proteobacteria (**Table S4**) and were chosen because we aimed to mutagenize several target bacteria from the Proteobacteria as a proof of concept. To construct this pool, we assembled each of these 10 designs separately, with random DNA barcodes. We then picked 10 colonies for each design and sequenced their DNA barcodes using BarSeq, and finally pooled all 100 clones together. Since each component part has been sequenced and archived, we know the absolute sequence information for every individual vector in the *mariner*-Kan magic pool, including its unique DNA barcode. We then transformed the magic pool into the *E. coli* conjugation donor strain WM3064 and performed BarSeq. BarSeq analysis showed that at least 9 out of the 10 DNA barcodes for each vector design were present in the conjugation strain culture, with an even distribution among the population (data not shown). We designed and constructed a Tn*5*-Kan magic pool in the same way as described above for the *mariner* transposon, this time based on the Tn5 RB-TnSeq transposon delivery vector pKMW7 (13). Again, we included 2 variants of the kanamycin resistance promoter (part2) and 5 variants of the transposase promoter (part5) (**Table S5**). All of the part5 transposase promoters were also from predicted essential genes from Proteobacteria genomes (**Table S5**).

We tested the two kanamycin magic pools against three environmental bacteria, *Sphingobium sp*. GW456-12-10-14-TSB1 (“Sphingo3”), *Sphingopyxis sp*. GW247-27LB (“Sphingo4”), and *Brevundimonas sp*. GW460-12-10-14-LB2 (“Brev2”). These bacteria were isolated from groundwater samples collected from the Oak Ridge National Laboratory Field Research Center (ORNL-FRC; see Materials and Methods) and each is sensitive to kanamycin. Because the *mariner*-Kan magic pool had higher mutagenesis efficiency with all three bacteria, we describe only the results for the *mariner* magic pool. We pooled about 5,000 Kan^R^ colonies from each conjugation, and TnSeq was performed on these preliminary mutant libraries. TnSeq maps both the transposon insertion site and its associated DNA barcode sequence (13). Because each DNA barcode has been previously associated with a specific transposon vector design, we can link each transposon insertion event back to a specific vector in the magic pool (**Figure 1**).

For each of the three of the preliminary mutant libraries from the *mariner*-Kan magic pool, DNA barcodes representing most of the 10 vector designs were observed, but their abundances varied (**Figure 2**). For Brev2, all 10 different designs produced transposon mutants, but pTGG36 through pTGG40 had higher efficiency compared to pTGG31 through pTGG35. pTGG31 to pTGG35 use the original Kan^R^ promoter from pKMW3 as part2, while pTGG36 to pTGG40 use the Amp^R^ promoter from pMarA (17). Similarly, in Sphingo3, the Amp^R^ promoter from pMarA as part2 was preferred, with pTGG37 being the most efficient vector (**Figure 2**). In Sphingo4, both part2 promoters had similar efficiency, with pTGG32 the most efficient vector overall (**Figure 2**). To verify the magic pool mutagenesis results, we constructed several individual transposon vectors and compared their mutagenesis efficiency in isolation against Brev2 and Sphingo3. In Brev2, we found that pTGG39 resulted more than 5 times the number of Kan^R^ mutants compared to pTGG32 or pTGG34. In Sphingo3, we found that pTTG37 produced at least 10 times the number of Kan^R^ mutants relative to pTGG34 or pTGG35. Therefore, for both Brev2 and Sphingo3, the magic pool results reflect the mutagenesis efficiency of the individual transposon vectors within the library.

**Figure 2.**
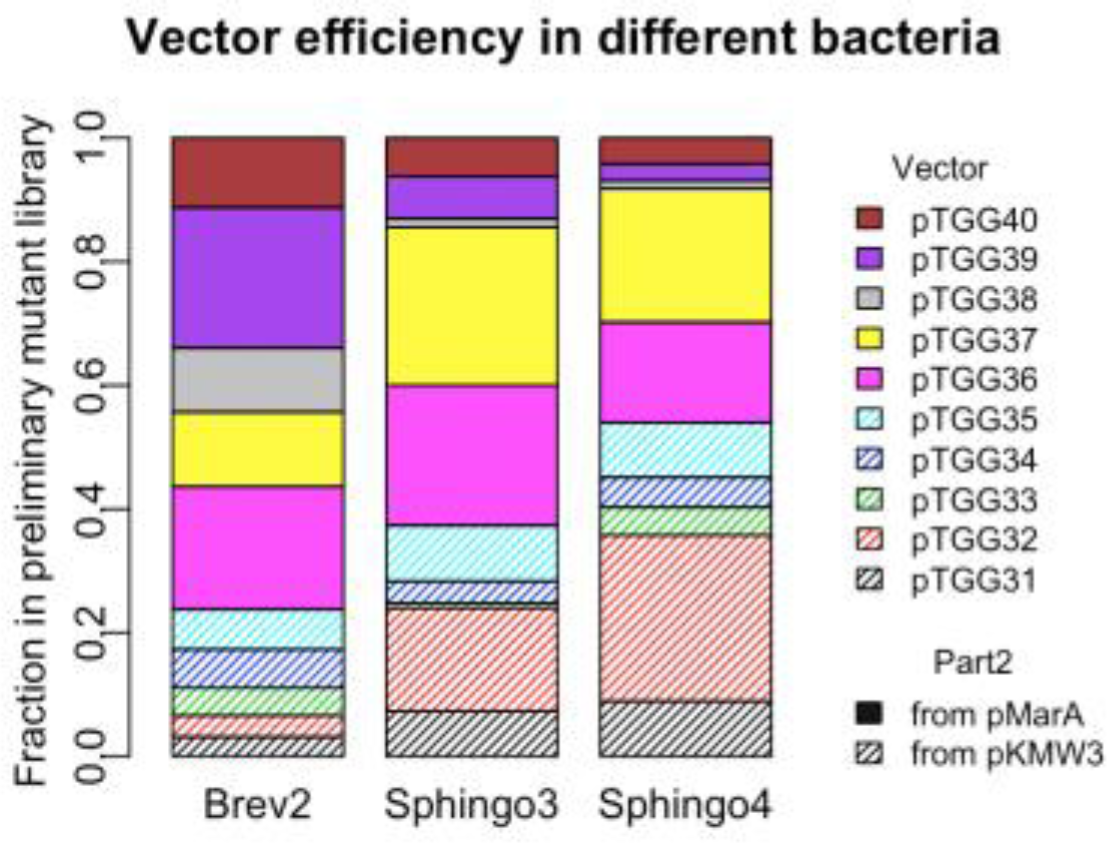
Mutagenesis efficiency of different *mariner* kanamycin vectors in magic pool. Based on barcode frequency associated with each vector in the *mariner*-kan magic pool, we show the fraction of unique insertions that map to each of the 10 transposon vector designs in the preliminary mutant library generated with the *mariner*-Kan magic pool. For a description of all of the parts in each vector, see **Table S4**. Brev2 = *Brevundimonas sp*. GW460-12-10-14-LB2; Sphingo3 = *Sphingobium sp*. GW456-12-10-14-TSB1; Sphingo4 = *Sphingopyxis sp*. GW247-27LB.

For the vectors in the magic pool that gave a sufficient number of insertions, we also examined the insertion events themselves (**Table S6**). First, we asked whether the insertions from that vector tended to be oriented with the antibiotic selection marker on the same strand as the disrupted gene. If they were, this could be a sign that the promoter for the resistance marker is not strong enough. We also asked if the insertions for that vector had an even number of insertions per gene, which is important for achieving good coverage. For example, in Brev2, pTGG36:pTGG40 all had decent efficiency with acceptable bias, and pTGG39 was the one with highest number of insertion locations and the lowest gene bias. So we chose pTGG39 as the best vector for Brev2. For similar reasons, we chose pTGG37 as the best vector for Sphingo3 (**Table S6**). Another issue was that in Sphingo4, pTGG32 had good efficiency but also had high read bias (mean reads per insertion location divided by median reads per insertion location = 327). In other words, some insertions gave far more reads than others. The high read bias could be a sign of selection for the loss of some gene (although it is not clear why this would occur only with this vector), an artifact of TnSeq, or some other issue. In any case, for Sphingo4, we would recommend using pTGG36 and/or pTGG37, which also had good efficiency but did not have the high read bias (**Table S6**).

To make full randomly barcoded transposon mutant libraries in Brev2 and Sphingo3, we first reassembled the individual vectors pTGG37 and pTGG39 using Golden Gate assembly with the archived parts vectors. In contrast to the magic pool construction, we used a part4 that was not yet barcoded. We then DNA barcoded (with a random 20-nucleotide sequence) the sequence-verified vectors using a second Golden Gate assembly-compatible enzyme (BsmBI; see Materials and Methods). We used two rounds of Golden Gate assembly, one to make the final vector, and a second to fully barcode it, to ensure high diversity of the barcodes for the RB-TnSeq workflow. We constructed two barcoded vectors, pTGG39_NN1 for Brev2, and pTGG37_NN1 for Sphingo3. As determined by BarSeq, the estimated barcode diversity in both barcoded vector libraries was greater than 10 million unique DNA barcodes (Materials and Methods), which is similar in diversity to our original RB-TnSeq vectors (13). Using these new randomly barcoded transposon vectors, we constructed whole-genome transposon mutant libraries in Brev2 and Sphingo3. We performed TnSeq to characterize each mutant library by mapping the insertion location and its associated random DNA barcode, as previously described (13). The Brev2 mutant library (Brev2_ML6) contains 166,981 mapped insertions and the Sphingo3 mutant library (Sphingo3_ML4) contains 275,037 mapped insertions (See **Table 1** for full details on the mutant libraries).

**Table 1.** Summary statistics for full mutant libraries.

| Category | *Sphingobium* sp.<br>GW456-12-10-14-TSB1 | *Brevundimonas*<br>sp. GW460-12-10-14-LB2 | *Echinicola*<br>*vietnamensis*<br>DSM17526 | *Pontibacter*<br>*actinarius*<br>DSM19842 | *Pedobacter* sp.<br>GW460-11-11-14-LB5 |
| --- | --- | --- | --- | --- | --- |
| Summary of mutant libraries |  |  |  |  |  |
| Mutant library name | Sphingo3_ML4 | Brev2_ML6 | Cola_ML5 | Ponti_ML7 | Pedo557_ML3 |
| Transposon | *mariner* vector<br>(pTGG37_NN1) | *mariner* vector<br>(pTGG39_NN1) | *mariner* vector<br>(pTGG43_NN2) | *mariner* vector<br>(pTGG43_NN2) | *mariner* vector<br>(pTGG43_NN2) |
| Method of delivery | conjugation | conjugation | conjugation | conjugation | conjugation |
| No. of strains with unique barcodes<sup>a</sup> | 275037 | 166981 | 278255 | 190948 | 190892 |
| % of barcodes with intact vectors<sup>b</sup> | < 0.1% | < 0.1% | < 0.1% | < 0.1% | < 0.1% |
| Protein coding genes |  |  |  |  |  |
| Total no. | 4903 | 3210 | 4625 | 4546 | 5229 |
| No. with central insertions<sup>c</sup> | 3943 | 2584 | 4090 | 4010 | 4670 |
| Median no. of strains per gene<sup>d</sup> | 30 | 27 | 32 | 19 | 21 |
| Insertion bias |  |  |  |  |  |
| % toward coding strand of genes | 50.6 | 49.9 | 48.2 | 49.6 | 48.8 |
| Mean/median no. of reads per gene | 1.60 | 1.68 | 1.49 | 1.42 | 1.40 |
| BarSeq statistics |  |  |  |  |  |
| % of reads with barcode | 98 | 98 | 98 | 98 | 98 |
| % of barcodes that map to a strain<sup>e</sup> | 60 | 75 | 61 | 73 | 80 |
| No. of reads per million for the median gene<sup>e</sup> | 65 | 109 | 66 | 69 | 83 |

<sup>a</sup> Only strains with insertions in the genome are included.<sup>b</sup> May represent integration events of entire transposon plasmid into genome.<sup>c</sup> with transposon lies within the central 10 to 90% of a gene.<sup>d</sup> Includes only protein coding genes with at least one central insertion.<sup>e</sup> Computed with time-zero samples.

To assess the utility of these mutant libraries for whole-genome mutant fitness assays using BarSeq, we carried out carbon utilization experiments for each bacterium. We grew each mutant library in defined minimal medium with glucose as the sole carbon source (see Materials and Methods). We define strain fitness as the log_2_ ratio of the strain’s abundance at the end versus the start of the assay, and we define gene fitness as the average of the strain fitness for insertions within the central 10-90% of the gene (13). As a metric for biological consistency, we first compared the log_2_ fitness values calculated between the two halves of each gene (13) (**Figure 3**). Specifically, we divided the mutants that have centrally-located insertions in the same gene into two halves, first half and second half, based on the transposon insertion location. Theoretically, the fitness value of the two halves should be consistent for most genes, though there are rare cases where the functional domain(s) is only located in one of the halves. Overall, we found a strong correlation between the first and second half gene fitness values for each organism, which demonstrates the internal consistency of the fitness values (**Figure 3**). As a second measure of biological consistency, we examined the genes that had fitness defects in defined media with glucose and found that many of them were predicted auxotrophic genes (TIGRFAMs with top-level role “Amino acid biosynthesis” (18)) or genes involved in glucose metabolism (**Table S7**). For example, in Sphingo3, we found that phosphogluconate dehydratase and a predicted mannose transporter, which we believe is more likely a glucose transporter based on homology (19), were both important for fitness. In Brev2, we identified glucose-6-phosphate isomerase as important for fitness in defined glucose-containing media. Interestingly, in both organisms, we also found that mutants of a capsule biosynthesis gene cluster have a significant growth disadvantage in defined media (**Figure 3, Table S7**).

**Figure 3.**
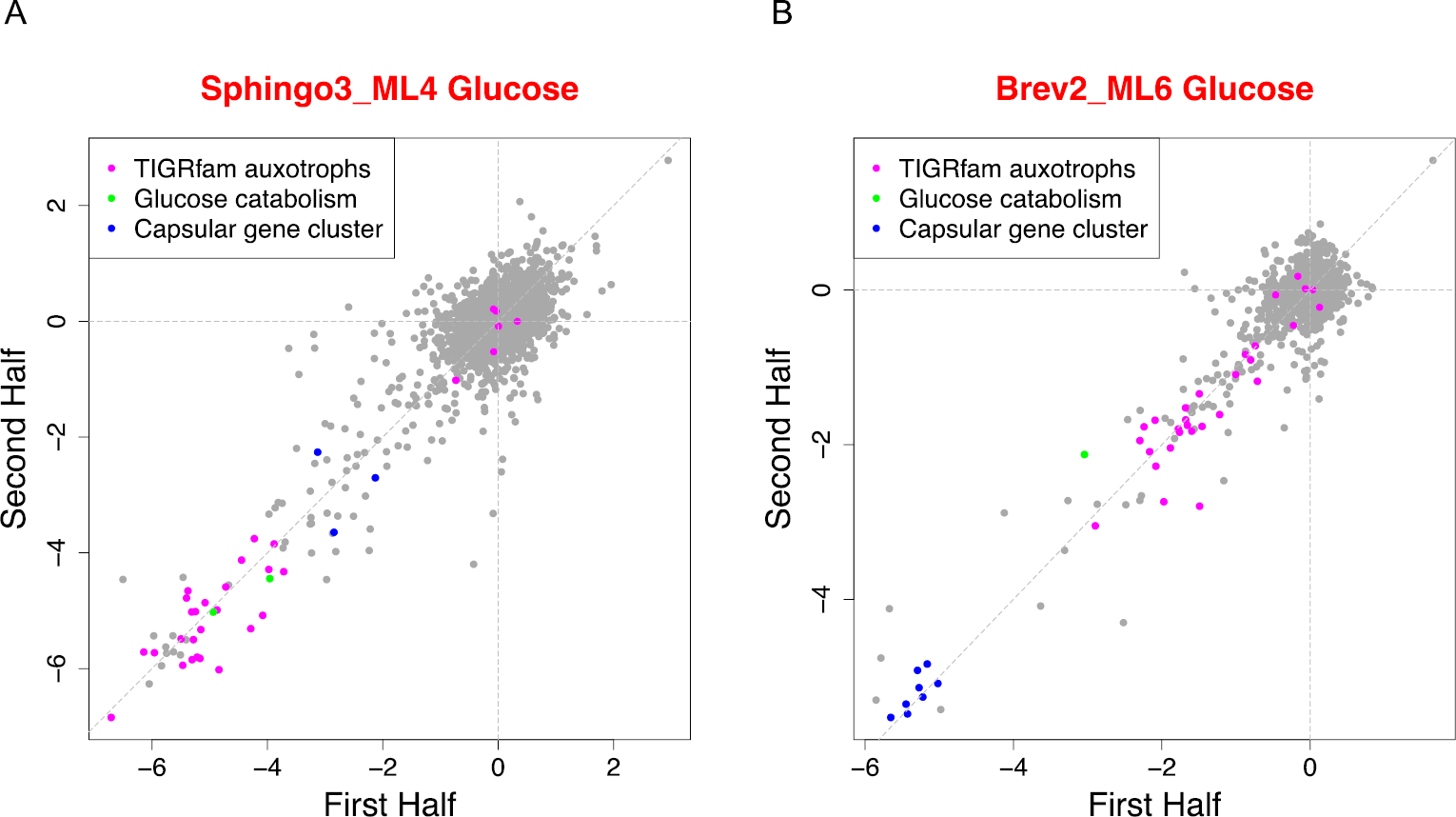
Mutant fitness data for two Proteobacteria. Fitness values for each gene as calculated using transposon insertions in the first versus second half (13). Each fitness value is a log_2_ ratio comparing the abundance of barcode abundance before and after growth selection. Genes highlighted in green and blue are listed in **Table S7**. (A) The mutant library of *Sphingobium sp*. GW456-12-10-14-TSB1 (Sphingo3_ML4) grown in defined media with glucose as a carbon source. Highlighted genes are (B) The mutant library of *Brevundimonas sp*. GW460-12-10-14-LB2 (Brev2_ML6) grown in defined media with glucose and a mixture of all 20 amino acids (see Materials and Methods). TIGRFAM auxotrophs are predicted amino acid biosynthesis genes (18).

### Large magic pools with erythromycin as the antibiotic selection marker

To extend our approach, we constructed two magic pools using erythromycin (Erm) as the selective antibiotic, one with Tn5 and one with *mariner*. To enable the mutagenesis of diverse bacteria, these magic pools were more complex than the kanamycin magic pools and they include promoters from a wider range of bacteria. For the Tn*5*-Erm magic pool, we used 10 variants of the antibiotic resistance gene promoter (part2), 5 variants of the Erm^R^ antibiotic resistance gene (part3), and 25 variants of the transposase promoter (part5), resulting in 1,250 possible combinations of parts (**Table S8**). For the *mariner*-Erm magic pool, we used the same 10 variants of part2 and the same 5 variants of part3, and 24 variants of part5, giving 1,200 possible vector combinations (**Table S9**). 12 of the part5 variants are used in both the *mariner* and Tn*5* Erm magic pools, while the remainder are unique to one magic pool. With these magic pools, we aimed to mutagenize members of the phylum *Bacteroidetes*, many of which are naturally resistant to kanamycin but sensitive to erythromycin. Therefore, among the part5 variants for Erm magic pools, 11 from Tn*5*-Erm and 12 from *mariner*-Erm are from members of the *Bacteroidetes* (****Table S8**; **Table S9****).

Due to the much higher number of possible transposon delivery vectors, we assembled and characterized the Erm magic pools in a different way than for the Kan magic pools. For Golden Gate assembly, we pre-mixed all the variants for a particular part with equal molar proportions, and then equal amounts of each part mixture were used for the final assembly (Materials and Methods). We pooled about 6,000 transformants for each Erm magic pool, representing ∼5 unique barcodes for each possible combination of parts. To characterize the Erm magic pools, we performed long read sequencing (PacBio) to link the DNA barcode to each of the parts on the same vector. In addition, we performed BarSeq to accurately assess the sequence of each DNA barcode in each magic pool. We were able to identify 618 different combinations of parts that were linked to at least one unique barcode in the Tn*5*-Erm magic pool and 638 different combinations in the *mariner*-Erm magic pool. Most of the individual parts were associated with at least one barcode and hence with at least one vector. However, we did not detect parts 2.9, 2.10, or 3.4 in any design for either Erm magic pool.

To test the two Erm magic pools, we selected three organisms from the phylum *Bacteroidetes: Pontibacter actiniarum* DSM19842 (“Ponti”) (20), *Echinicola vietnamensis* DSM17526 (“Cola”) (21), and *Pedobacter sp*. GW460-11-11-14-LB5 (“Pedo557”), another ORNL-FRC isolate. Both Cola and Ponti are naturally resistant to kanamycin and sensitive to erythromycin. We made a preliminary mutant library with *mariner*-Erm in all three bacteria. We also made preliminary mutant libraries with Tn*5*-Erm in Ponti and Cola. For Pedo557, the mutagenesis efficiency of the Tn*5*-Erm magic pool was too low to make a preliminary mutant library. To identify the best vector parts, we performed TnSeq on the preliminary mutant libraries to identify the DNA barcode and the transposon insertion location for each mutant in the pools.

For driving the expression of the antibiotic selection marker, p2.7 (the *dnaE* promoter of *Bacteroides thetaiotaomicron* strain VPI-5482) was the dominant part2 variant in both Cola and Ponti (**Figure 4**). It was also one of the two most abundant part2 variants in Pedo557, with the other one being a DNA ligase promoter from *Bacteroides thetaiotaomicron* (p2.8).

**Figure 4.**
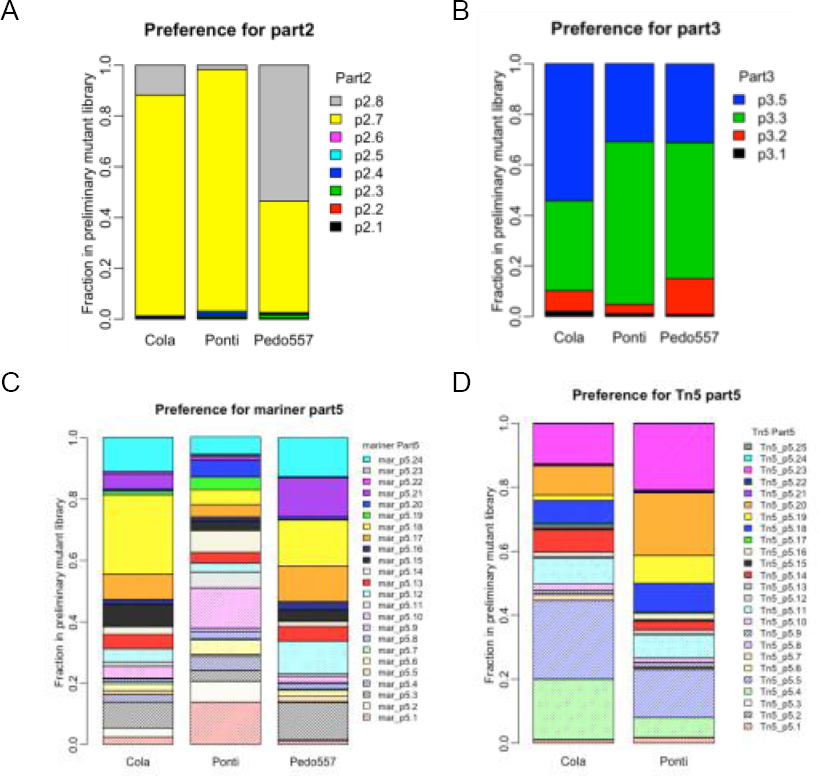
Preference for part variants in the Erm magic pools. (A) As determined by DNA barcodes identified by TnSeq, the fraction of each of 8 part2 variants in the combined *mariner* and Tn5 preliminary mutant libraries for *Echinicola vietnamensis* ("Cola”), *Pontibacter actiniarum* (“Ponti”), and *Pedobacter sp*. GW460-11-11-14-LB5 (“Pedo557”). (B) The fraction of each of the 4 part3 variants in the combined *mariner* and Tn5 preliminary mutant libraries for each bacterium. (C) The fraction of each of the 24 *mariner* part5 variants in Cola, Ponti, and Pedo557. (D) The fraction of each of the 25 Tn5 part5 variants in Cola and Ponti.

Among all 4 variants of the open reading frame of the 23S rRNA methytransferase conferring erythromycin resistance as part3, the two most successful variants were p3.3 and p3.5 (**Figure 4**). p3.3 is the *ermBP* gene from *Clostridium perfringens* (22). p3.5 is the *ermC* gene from plasmid pE194, which was originally isolated from *Staphylococcus aureus* (23). Part p3.2 is derived from a plasmid for genetic engineering of Clostridium (24) and differs by only 5 amino acids from p3.5, but it was much less efficient than p3.5. The other variant p3.1, the *ermF* gene from *Bacteroides fragilis* (25), was detected rarely. The less successful variants of part3 might not express well in our three target bacteria, possibly due to codon preference and/or negative interactions between the ORF and the ribosome binding site (26).

For the promoter driving expression of the transposase (part5), almost all of the variants were detected in the preliminary mutant libraries (**Figure 4**). For Cola, with the *mariner* magic pool, the *Dxs* promoter from *Belliella baltica* (mar_p5.18) was about 25% of the part5 variants detected in the *mariner* mutant library, followed by the *rfaG* promoter from *Pontibacter actiniarum* (mar_p5.24) with 11.20% abundance. With the Tn5 magic pool, the three most abundant variants for Cola (>10%) were the *dnaE* promoter from *Prevotella multisaccharivorax* (Tn5_p5.4), the *kdsB* promoter from *Prevotella ruminicola* (Tn5_p5.5), and the *wecE* promoter from *Echinicola vietnamensis* (Tn5_p5.23). For Ponti, with the *mariner* magic pool, the *plsB* promoter from *Delftia sp*. GW456-R20 (mar_p5.1) and the *dnaE* promoter from *Collinsella stercoris* (mar_p5.10) were the most abundant (∼13%) (**Figure 4**). With the Tn5 magic pool, 3 promoters had abundance over 15% in the Ponti preliminary mutant library, two that were also abundant with Cola (Tn5_p5.5 and Tn5_p5.23) and Tn5_p5.20 (the *kdsB* promoter from *Pontibacter actiniarum)*. In the preliminary *mariner* mutant library of Pedo557, 6 part5 variants were observed at higher abundance, including two that were abundant in Cola, parts mar_p5.18 and mar_p5.24 (**Figure 4)**.

### Full mutant libraries in 3 Bacteroidetes

Because the *mariner* magic pool gave more transformants than Tn5 with our test conjugation in all 3 bacteria, we constructed the final transposon delivery vectors based on the *mariner* magic pool. All three bacteria showed a common preference for variants p2.7 and p3.3. To choose which part5 to use, we considered the bias of the vectors containing both p2.7 (the *dnaE* promoter from *Bacteroides thetaiotaomicron)* and p3.3 (the *ermBP* gene from *Clostridium perfringens)*. In all three bacteria, several variants of part5 seem to be functional without significant bias (**Table S10**). We chose to use mar_p5.18 (the *Dxs* promoter from *Belliella baltica)* as the transposase promoter for all three bacteria even though this part variant was not the most abundant for Ponti and Pedo557. However, we expected that it would be sufficient for full mutant library construction in both of these bacteria as its abundance in the magic pool data was reasonably high (>4% for both) (**Figure 4**). Based on this design, we assembled and DNA barcoded the *mariner* transposon delivery vector pTGG43_NN2 using p2.7, p3.3 and mar_p5.18 as the part2, part3 and part5, respectively. We successfully made 3 *mariner* transposon insertion mutant libraries using pTGG43_NN2: Cola_ML5, Ponti_ML7, and Pedo557_ML3. We performed TnSeq to characterize each mutant library by linking the random DNA barcode to its transposon insertion site. Each of the three mutant libraries contains at least 190,000 mapped insertions (**Table 1**).

We performed genome-wide mutant fitness assays in defined minimal media to confirm that we could generate biological meaningful data for each of the three mutant libraries. As shown in **Figure 5**, in both Cola and Pedo557, many predicted auxotrophs are required for optimal fitness in defined media. Further, these results are very consistent whether the gene fitness values are computed from insertions in the first versus second half of each gene (**Figure 5**). Glucose-6-phosphate isomerase from Cola and glucose-6-phosphate 1-dehydrogenase mutants from Pedo557 were both important for fitness in glucose-containing media, as expected (**Figure 5; Table S7**). We were unable to grow Ponti in a defined minimal medium supplemented with a single carbon source, but it grows well when supplemented with a mixture of the 20 standard amino acids. Therefore, most predicted Ponti auxtrophs display no fitness defect in this experiment. Rather, we found that a polysaccharide synthesis gene cluster was important for fitness in defined media with a mixture of the 20 amino acids (**Figure 5; Table S7**).

**Figure 5.**
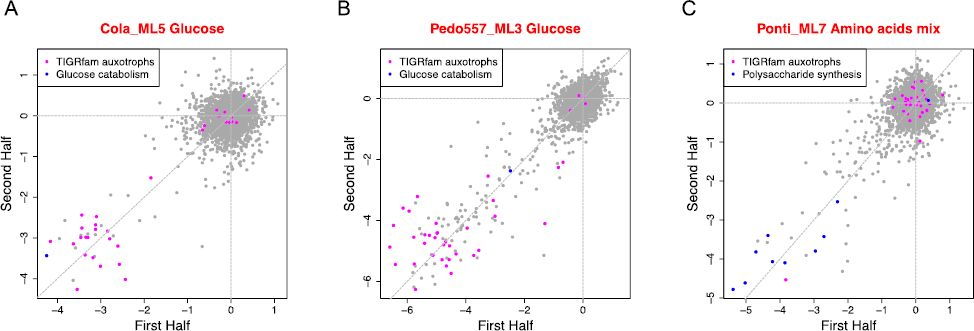
Genome-wide mutant fitness data for three *Bacteroidetes* strains. Fitness values for each gene as calculated using transposon insertions in the first versus second half (13). Each fitness value is a log_2_ ratio comparing the abundance of barcode abundance before and after growth selection. Genes highlighted in blue are listed in **Table S7**. (A) The mutant library of *Echinicola vietnamensis* (Cola_ML5) grown in defined minimal medium with glucose as the sole carbon source. (B) The mutant library of *Pedobactersp*. GW460-11-11-14-LB5 (Pedo557_ML3) grown in defined minimal medium with glucose as the sole carbon source. (C) The mutant library of *Pontibacteractiniarum* (Ponti_ML7) grown in a defined minimal media with a mixture of the 20 amino acids (see Materials and Methods). TIGRFAM auxotrophs are predicted amino acid biosynthesis genes (18).

## Discussion

In this study we present a strategy for testing the efficacy of hundreds of transposon vectors in parallel against a target bacterium. The purpose of this work was to accelerate the identification of a working construct for RB-TnSeq (13), rather than laboriously testing transposon vectors individually. Our parts-based strategy involves the construction of a mixture of different transposon delivery vectors that can be simultaneously characterized by DNA sequencing due to the presence of a random DNA barcode that is included in each construct during Golden Gate assembly. A small preliminary mutant library containing a few thousand mutants and one round of TnSeq was sufficient to identify an effective vector for constructing the full mutant library. Once the ideal vector was chosen, it only takes about 2 weeks to reassemble the vector, fully barcode the vector with another round of Golden Gate assembly, and to make the full barcoded mutant library. This workflow is streamlined because all individual parts are archived for rapid vector reconstruction and because barcoding the final vector by Golden Gate assembly is easier than our original strategy of classic restriction enzyme-based cloning (13).

We found that the selectable marker and the promoter driving it were more important than which promoter was used to drive transposase expression. This suggests that high expression of the selectable marker in the transposon is critical for unbiased transposon mutagenesis. We speculate that the expression level of the transposase could vary to some extent in different host cells without (apparently) having a dramatic effect on the frequency of transposition, and without affecting the bias of the final library. In contrast, weak expression of the selectable marker could lead to bias (i.e, insertions on the sense strand of highly-expressed genes have a growth advantage because they have higher expression of the marker).

Although we only tested our four magic pools against five genera of bacteria, we expect that a wide range of Gram-negative bacteria could be mutagenized by at least one of the vectors in these pools. Furthermore, the magic pools could be easily expanded by incorporating more part variants, such as other selectable markers or additional promoters to drive the expression of the selectable marker. With the magic pools and RB-TnSeq, it should be feasible to collect extensive genome-wide mutant fitness data across dozens of conditions for diverse bacteria.

## Materials and Methods

### Bacterial strains

The bacterial strains used in this study are listed in **Table S1**. Competent cells of *Escherichia coli* TransforMax™ EC100D™ pir+ cloning strain and the WM3064 conjugation donor strain were purchased from Lucigen (www.Lucigen.com). *E. coli* 10-beta competent cells were purchased from New England Biolabs. The wild-type strains of *Pontibacter actiniarum* DSM 19842 (“Ponti”) and *Echinicola vietnamensis* DSM 17526 ("Cola”) were purchased from DSMZ (www.dsmz.de). *Sphingobium sp*. GW456-12-10-14-TSB1 (“Sphingo3”), *Sphingopyxis sp*. GW247-27LB (“Sphingo4”), *Brevundimonas sp*. GW460-12-10-14-LB2 (“Brev2”), and *Pedobacter sp*. GW460-11-11-14-LB5 (“Pedo557”) were isolated from the Oak Ridge National Laboratory field research center (ORNL-FRC; https://public.ornl.gov/orifc/orfrc3site.cfm). Sphingo3 was isolated on 10^-1^ diluted tryptic soy agar, Sphingo4 on LB agar, and Brev2 and Pedo557 on 10^-1^ diluted LB agar. All microbes were isolated aerobically at 30°C.

### Growth conditions

All growth media was purchased from BD and other chemical compounds were purchased from Sigma-Aldrich. *E. coli* was grown in Luria-Bertani (LB) medium at 37°C supplemented with antibiotics as needed: 50 μg/ml carbenicillin or 20 μg/ml chloramphenicol. For culturing WM3064, a final concentration of 300 μM diaminopimelic acid (DAP) was supplemented in LB. We grew Sphingo3, Sphingo4, Brev2 and Pedo557 in R2A medium at 30°C. We grew Cola and Ponti in BD marine broth 2216 at 30°C. For mutant selection and culturing, we supplemented the medium with antibiotics at the following concentration: 25 μg/ml kanamycin for Sphingo3 and Sphingo4 mutants, 100 μg/ml kanamycin for Brev2 mutants, and 25 μg/ml erythromycin for Cola, Ponti and Pedo557 mutants. For growth on solid media, we used 1.5% (w/v) agar.

### Genome sequencing

We sequenced the genomes of *Brevundimonas sp*. GW460-12-10-14-LB2, *Pontibacter actiniarum, Pedobacter sp*. GW460-11-11-14-LB5, and *Sphingobium sp*. GW456-12-10-14-TSB1 with a combination of long reads from PacBio and short reads from Illumina. We used RS_HGAP_Assembly.3 (27) in the SMRT Portal to assemble the PacBio reads, circulator (28) to circularize any complete contigs, and pilon (29) to correct local errors using Illumina reads. We sequenced the genome of *Sphingopyxis sp*. GW247-27LB using Illumina and the A5 assembler (30). For *Pontibacter actiniarum*, we downloaded preexisting short Illumina reads from the JGI; all of the other genome sequencing data was generated for this project.

These genome sequences have been deposited into Genbank. *Brevundimonas sp*. GW460-12-10-14-LB2 is accession CP015511. *Pontibacter actiniarum* KMM 6156 (DSM 19842) is accession CP021235-CP021236. *Sphingobium sp*. GW456-12-10-14-TSB1 is accession NGUN00000000. *Pedobacter sp*. GW460-11-11-14-LB5 is accession CP021237. *Sphingopyxis sp*. GW247-27LB is accession NIWD00000000.

We used RAST (31) to annotate protein-coding genes. To identify genes that are expected to be involved in amino acid biosynthesis, we assigned genes to TIGRFAMs 15.0 (18) using HMMer 3.1b1 (32) and used all TIGRFAMs with the top-level role “Amino acid biosynthesis”.

### Construction of the parts vectors

The plasmids used in this study are listed in **Table S2** and the oligonucleotides/gBlocks in **Table S3**. All oligonucleotides and gBlocks were ordered from IDT (www.idtdna.com). Unless noted otherwise, all PCR reactions were carried out using the Q5 hot start DNA polymerase from NEB. DNA segments were cleaned up and/or concentrated using the DNA Clean & Concentrator kit (Zymo Research). Gibson assembly reactions were carried out using the Gibson Assembly Master Mix from NEB. Golden Gate assembly enzymes Esp3I (BsmBI) and BpiI (BbsI) were purchased from Thermo Fisher Scientific. T4 DNA ligase and buffer were purchased from NEB. Plasmid isolation was done using the QIAprep Spin Miniprep Kit (Qiagen).

Our transposon vector construction and DNA barcoding strategies both utilize Golden Gate assembly (15, 16). For transposon vector assembly, we use the Golden Gate compatible enzyme BbsI (**Figure 6**). For random DNA barcoding of transposon delivery vectors, we use a second Golden Gate compatible enzyme, BsmBI. Therefore, our approach requires that the individual parts not have recognition sequences for BbsI (5’ GAAGAC) or BsmBI (5’ CGTCTC). We started by constructing a universal part-holding vector pJW52. pJW52 is derived from pML967, a vector with a colE1 origin of replication, the *cat* gene conferring resistance to chloramphenicol, and GFP. To make pJW52, we first performed site-directed mutagenesis with oligonucleotides ofeba243 and ofeba244 (using the NEB Q5 site-directed mutagenesis kit) to remove a BbsI site present in pML967. We then performed a second of site-directed mutagenesis with oligos ofeba445 and ofeba446 to remove a BsmBI site from pML967. See **Table S2** for construction details for all plasmids used in this study.

**Figure 6.**
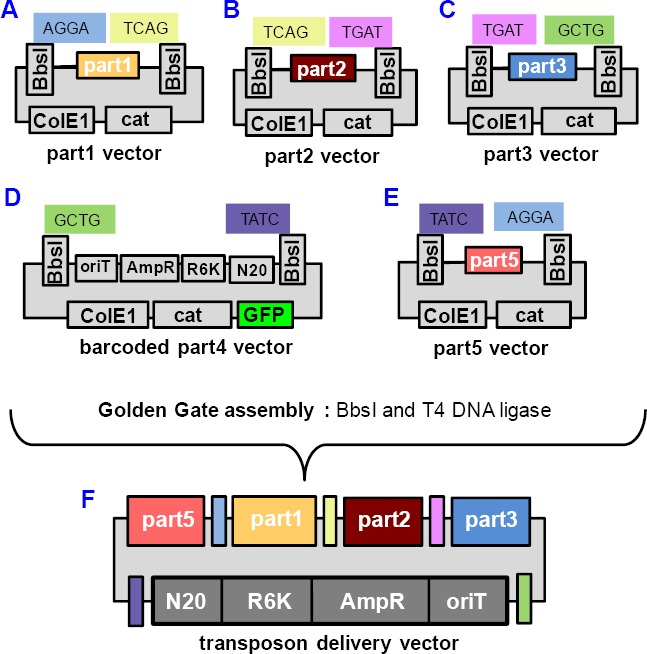
Golden Gate assembly of transposon delivery vectors from part vectors. (A-E) The 5 part vectors that are used for Golden Gate assembly of the magic pool transposon delivery vectors: part1 vector, part2 vector, part3 vector, barcoded part4 vector and part5 vector (not drawn to scale). We show the sequences of the 4 nucleotide overhangs for Golden Gate assembly. (F) The transposon delivery vector with DNA barcode. ColE1: replication origin colE1; cat: chloramphenicol resistance gene; GFP: green fluorescent protein; oriT: origin of transfer; ampR: beta-lactam resistance gene; R6K: conditional replication origin; N20: random 20 nucleotide DNA barcode.

To construct the Tn5 part1 vector pHLL212, we first constructed the intermediate vector pJW8 by moving most of the Tn5 transposase, the R6K origin of replication, and one IR from pKMW7 (13) into pML967. To make pHLL212, we removed the R6K origin of replication from pJW8 by PCR and self-ligation. The mariner part1 vector pHLL213 was constructed in a similar way, we first make the intermediate vector pJW20 by moving most of the *mariner* transposase, the R6K origin of replication, and one IR from pKMW3 into pML967. We then removed the R6K origin of replication from pJW20 by PCR and self-ligation to make pHLL213.

To construct the non-barcoded Tn5 part4 vector pHLL214, we first made the intermediate vector pJW54 by cloning a gBlock (gfeba443) and a portion of pUC19 into pJW52, linearized with ofeba247 and ofeba248. To make pHLL214, we then sewed together two PCR products by Gibson assembly, oHL557-oHL558 amplified pJW54 and oHL561-oHL562 amplified pJW20. The non-barcoded *mariner* part4 vector pHLL215 was constructed in a similar way (see **Table S2** for details). The part4 vectors contains the R6K origin of replication, the *bla* gene conferring resistance to carbenicillin for cloning, the conjugation origin of transfer, the second transposon IR, and a short region for randomly DNA barcoding the vector via Golden Gate assembly (see below). The part4 vectors are the only part vectors to contain GFP on the backbone. The part4 vector is designed in this way, such that after the Golden Gate assembly reaction, a correct transposon vector assembly will lose the GFP cassette, and the colonies would not appear green on plates. We use the number of green color colonies vs total number of all colonies as an indicator for the assembly efficiency.

To construct the two randomly DNA barcoded part4 vectors, pHLL214_NN1 and pHLL215_NN1, we first PCR amplified a short 46 bp oligo with 20 random base pairs in the middle (ofeba282) with oligos ofeba285 and ofeba286 using Phusion DNA polymerase (98°C 1 minute, 6 cycles of 98°C 10 seconds, 58°C 30 seconds, 72°C 60 seconds, followed by 72°C for 5 minutes). The barcode PCR products were purified with the DNA Clean & Concentrator kit (Zymo Research) and included in a Golden Gate assembly reaction with either pHLL214 or pHLL215. We performed the Golden Gate assembly reaction with 1000 ng of part4 plasmid (pHLL214 or pHLL215) and 40 ng of the barcode PCR product using BsmBI (Esp3I from Thermo Fisher) and T4 DNA ligase under the following cycling conditions: 10 cycles of 37°C for 5 minutes and 16°C for 10 minutes, followed by a final digestion step at 37°C for 30 minutes. We then purified the Golden Gate assembly reaction with the DNA Clean & Concentrator kit (Zymo Research) and digested the DNA overnight with BsmBI. We performed this additional digestion step to further eliminate non-barcoded vectors. We then purified the DNA again with the DNA Clean & Concentrator kit (Zymo Research) and transformed into *E. coli*.

The part2 (antibiotic marker promoter), part3 (ORF of the antibiotic selection marker), and part5 (transposase promoter and 5’ end of the transposase) parts were mostly cloned into the universal holding vector pJW52, amplified with ofeba134 and ofeba137 (**Table S2**). In some occasions, the sequence was slightly modified to eliminate the BbsI and BsmBI recognition site(s), and site mutagenesis was used when necessary. The part5 promoters were chosen from predicted essential genes in diverse bacteria. To do this, we made a list of 22 COGs that are expected to be essential and are moderately expressed in *Shewanella oneidensis* MR-1. If a genome of interest had just one member of the COG and it was the first gene of the operon, then we selected it as a candidate promoter. The part variants were mainly synthesized as gBlocks with the exception that a few were generated by PCR amplification using long overlapping oligos as the template (**Table S2**). All of these parts were generated by Gibson assembly and sequence-verified.

### Construction of the kanamycin magic pools

Since there were only two part2 variants and five part5 variants, with a total of 10 possible different constructs, we constructed those 10 designs separately (pTGG21-pTGG30 for the Tn5-kan magic pool, and pTGG31-pTGG40 for the *mariner*-kan magic pool, more details in **Table S4** and **Table S5**). The barcoded part4 vector was used for making the magic pools. BbsI was used for the Golden Gate assembly reaction (using the same protocol and cycling conditions described above for barcoding the part4 vectors), and the assembly product was transformed into *E.coli* Pir+ cloning strain. Then 10 GFP negative colonies from each transformation were randomly picked and the barcode region was sequenced to make sure that each had a unique barcode. For the kanamycin-based magic pools, since we constructed each design for each magic pool individually, we performed BarSeq to identify the barcodes used for each of them before we combined the 100 vectors to make the final kanamycin magic pools. Barcodes with at least 5 reads were confidently associated with that design.

### Construction of the erythromycin magic pools

We constructed the larger erythromycin magic pools using Golden Gate assembly with the same methods as for the kanamycin magic pools, except that we used more variants for some parts. Specifically, the *mariner*-Erm magic pool contains 10 variants of part2, 5 variants of part3 and 24 variants of part5, and 10, 5 and 25 variants for each part respectively, for the Tn*5*-Erm magic pool (**Table S8** and **Table S9**). We used 100 ng for each part mixture for the Golden Gate assembly reaction. For example, since we used 10 different variants of part2, the amount of each of the part2 variants was about 10 ng. We used the same Golden Gate cycling conditions described above for barcoding the part4 vectors except that we used BbsI to assemble the transposon vectors. We mixed ∼6000 colonies to make the *mariner*-Erm and Tn*5*-Erm magic pools.

For each of the erythromycin-based magic pools, which were far more complex, we used PacBio to sequence the entire mixture of the plasmids as well as barcode sequencing (BarSeq) with Illumina to identify the exact sequences of the barcodes that were present. To analyze the long reads from PacBio, we first obtained circular consensus reads using the “Reads for Insert” method in the SMRT portal. We then used BWA (33) to map these reads to the expected parts. We also used BWA to identify the “seq2” region that lies jus t downstream of the barcode. We then extracted the part of the consensus read that was expected to contain the barcode, plus two additional nucleotides on each side. We then matched the PacBio barcodes (which are very noisy) to a database of actual barcodes as determined from Illumina sequencing (BarSeq), using blastn. In the Illumina data, we assumed that the most abundant barcodes that account for 98% of the reads are genuine; the remaining barcodes were ignored as potential sequencing error. The BLAST database included 2 additional flanking nucleotides on each side. Any hit of at least 30 bits and on the correct strand was considered to be the correct barcode, unless there was another barcode within one bit. We linked this barcode to all of the parts that were identified. If a part was identified with low confidence (a mapping quality below 10), then that part was considered to be unknown. Additionally, some reads did not contain any instance of a part. Finally, given a list of reads that map a barcode to one or more parts, we took the combination of all these mappings to generate our definition of the magic pool. If a barcode mapped to more than one instance of a part, then that part was considered to be unknown.

For pHLL254 (the Tn*5*-Erm magic pool), we obtained 36,935 circular consensus reads from two cells of PacBio. Of these, 16,084 reads contained a putative barcode region, 13,355 reads had a barcode confidently identified, and 10,627 reads linked a barcode to at least one part. Overall, 4,766 barcodes in pHLL254 (of the 6,690 confident barcodes in the Illumina BarSeq data) were linked to at least one part that varies, 2,526 barcodes were linked to all three of those parts, and 618 different combinations of the three variable parts were linked to at least one barcode.

For pHLL255 (the *mariner*-Erm magic pool), we obtained 37,791 circular consensus reads with two cells of PacBio. Of these, 14,174 reads contained a putative barcode region, 11,177 reads had a barcode confidently identified, and 9,398 reads linked a barcode to at least one part. Overall, 4,639 barcodes in pHLL255 (of the 6,635 confident barcodes in the Illumina BarSeq data) were linked to at least one part that varies, 2,918 barcodes were linked to all three of those parts, and 638 different combinations of the three variable parts were linked to at least one barcode.

### Barcoding of individual vectors

Once we picked a suitable transposon delivery vector for a particular bacterium, we identified the specific variant for each part, and we then reassembled the vector using Golden Gate assembly as described above with BbsI, except that we used the non-barcoded version of part4. We then incorporated random DNA barcodes into the vector using the exact same strategy described above for barcoding the part4 vectors pHLL214 and pHLL215. The barcoded vector was then transformed into the *E. coli* Pir+ strain, with between 10-100 million transformants. We picked 20 colonies at random and sequenced the barcode region to give an estimation about the barcoding efficiency, which was over 90% in all cases. BarSeq analysis was performed to estimate the total unique barcodes in each barcoded transposon delivery vector. For example, for pTGG37_NN1, we had 2.55 million high-quality BarSeq reads, that is, every position of the barcode had a quality score of at least 30. This implies that less than 2% of these reads had any errors in the barcode. We observed 2.32 million different barcodes, of which 5,603 are one nucleotide different from another barcode and probably represent sequencing errors. We observed 2.12 million barcodes just once and 0.19 million barcodes twice. If we use Chao’s estimator and we (conservatively) reduce the number of distinct barcodes and singleton barcodes to account for 2% of them being potential sequencing errors, then we estimate that there are about 13.7 different million barcodes in pTGG37_NN1. We then transformed the barcoded vector library into the *E. coli* conjugation donor strain WM3064 and performed another round of BarSeq to confirm that the barcode diversity in the conjugation donor strain was comparable to the diversity in the Pir+ cloning strain.

### Transposon mutagenesis

We carried out transposon mutagenesis by conjugating the recipient cells with the *E. coli* donor cells carrying the barcoded transposon vectors (either a magic pool, or the fully barcoded single vectors) at 1:1 ratio. Specifically, for making the Pedo557 preliminary mutant library with the *mariner*-Erm magic pool, we first grew 10 mL of wild-type Pedo557 in R2A overnight at 30°C. The next morning, we recovered a 2 mL freezer stock of the *mariner*-Erm magic pool in the conjugation donor strain (AMD280) in 50 mL LB supplemented with carbenicillin and DAP, at 37°C. When the OD_600_ of the *E. coli* donor strain reached about 1, we harvested 3 OD_600_ units of the culture and washed the cells 3 times with fresh R2A supplemented with DAP. Then 3 OD_600_ units of the Pedo557 wild-type cells were harvested and mixed with the washed donor strain cells, then resuspended to the final volume of 60 μL with R2A liquid medium supplemented with DAP. The resuspension was spotted onto a MF-Millipore 0.45 μm Gravimetric Analysis Membrane Filter, and incubated overnight on an R2A agar plate supplemented with DAP at 30°C. The next day, the conjugation mixture was scrapped off from the membrane and resuspended into 2 mL fresh R2A medium supplemented with erythromycin (25 μg/ml for Pedo557), then we plated a series of dilutions onto R2A plates supplemented with erythromycin. The plates were incubated at 30°C for about 48 hours to let visible colonies develop. We then pooled ∼5000 colonies to construct the Pedo557 *mariner* magic pool preliminary mutant library. The other preliminary mutant libraries were constructed using a similar strategy, with only minor changes to the media used (see above). To construct the 5 full mutant libraries described in this study, we followed the same conjugation method described above, except that we pooled together over 100,000 colonies. In practice, we typically constructed multiple large mutant libraries for each bacterium, as colony counts may be misleading. For each large-scale mutant library, we performed a preliminary round of BarSeq to estimate the size of the mutant library using the same Chao estimator described above. We then proceeded with TnSeq for the mutant library with a predicted size of 100,000 to 300,000 mutant strains. For each full mutant library, we made multiple, single-use glycerol stocks of the library and extracted genomic DNA for TnSeq analysis. To map the genomic location of the transposon insertions and to link these insertions to their associated DNA barcode, we used the same TnSeq protocol that we described previously (13).

### Genome-wide mutant fitness assays

We performed genome-wide mutant fitness assays with BarSeq as described (13). Briefly, we thawed an aliquot of the full transposon mutant library, inoculated the entire aliquot into 25 mL of media (R2A or marine broth depending on the bacterium) with the appropriate antibiotic (either kanamycin or erythromycin), and grew the library at 30°C until the cells reached mid-log phase. We then collected cell pellets (the “time-zero” samples) and used the remainder of the cells to set up competitive growth assays in defined minimal media. We washed the cells 3 times with a 2X concentrated defined media minus a carbon source. We then inoculated cultures for fitness assays in 5 mL total volume (in 15 mL culture tubes) by mixing 2.5 mL 2X concentrated carbon source with 2.5 mL 2X concentrated media minus carbon containing cells at an OD_600_ of 0.04 to give a final OD_600_ of 0.02 in 1X media with 1X carbon source. As the baseline growth media, we used ShewMM_noCarbon for Brev2 and Sphingo3, RCH2_defined_noCarbon for Pedo557, and DinoMM_noCarbon_HighNutrient for Cola and Ponti. The components for each of these growth media are described (13). We used a 20 mM final concentration of D-glucose as the carbon source for all bacteria except Ponti. For Ponti, we used a mixture of the 20 standard amino acids mixed in equal proportions such that the final concentration of each amino acid was 0.25 mM. We also added the amino acid mixture to the Brev2 experiment (each amino acid at 0.05 mM), as we found that this stimulated the growth of this bacterium. After the mutant library reached stationary phase (after 1 to 3 days of growth, depending on the mutant library), we collected cell pellets (the “condition” sample). We extracted genomic DNA from the time-zero and conditions samples, PCR amplified the DNA barcodes, and sequenced the barcodes using Illumina (BarSeq). We performed BarSeq as previously described (13), except for a change to our common P1 oligo design. We used an equimolar mixture of P1 oligos with 2-5 random N’s to “phase” our amplicons and to support sequencing with the Illumina HiSeq4000 (**Table S3**). Gene fitness values were calculated as described (13).

## Data availability

For data describing the magic pools, the full mutant libraries, and the mutant fitness assays, as well as source code, see http://genomics.lbl.gov/supplemental/magicpools/.

## Acknowledgements

This material by ENIGMA – Ecosystems and Networks Integrated with Genes and Molecular Assemblies (http://engima.lbl.gov), a Scientific Focus Area Program at Lawrence Berkeley National Laboratory is based upon work supported by the U.S. Department of Energy, Office of Science, Office of Biological & Environmental Research under contract number DE-AC02-05CH11231.

## Supplementary Info

**Table S1**: Strains used in study

**Table S2**: Plasmids used in study

**Table S3**: Oligonucleotides and gBlocks used in study

**Table S4**: The *mariner*-Kan magic pool

**Table S5**: The Tn5-Kan magic pool

**Table S6**: Kan magic pool analysis

**Table S7**: Select genes with fitness defects

**Table S8**: The Tn*5*-Erm magic pool

**Table S9**: The *mariner*-Erm magic pool

